# How to better conserve with genetic data: origin and population structure of Brazilian locally adapted hair sheep (*Ovis aries*) breeds

**DOI:** 10.1101/815217

**Authors:** Tiago do Prado Paim, Samuel Rezende Paiva, Natália Martins de Toledo, Michel Beleza Yamaghishi, Paulo Luiz Souza Carneiro, Olivardo Facó, Adriana Mello de Araújo, Hymerson Costa Azevedo, Alexandre Rodrigues Caetano, Concepta McManus

**Affiliations:** Faculdade de Agronomia e Medicina Veterinária, Campus Darcy Ribeiro, Universidade de Brasília, Brasília, Distrito Federal, Brazil.; Instituto Federal de Educação, Ciência e Tecnologia Goiano, Iporá, Goiás, Brazil.; Embrapa Recursos Genéticos e Biotecnologia, Brasília, Distrito Federal, Brazil.; Embrapa Informática Agropecuária, Campinas, São Paulo, Brazil.; Departamento de Ciências Biológicas, Universidade Estadual do Sudoeste da Bahia, Jequié, Bahia, Brazil.; Embrapa Caprinos e Ovinos, Sobral, Ceará, Brazil.; Embrapa Meio-Norte, Teresina, Piaui, Brazil.; Embrapa Tabuleiros Costeiros, Aracaju, Sergipe, Brazil.; Instituto de Biologia, Universidade de Brasília, Brasília, Distrito Federal, Brazil.

## Abstract

Brazilian hair sheep constitute a genetic diversity hotspot of sheep breeds. These locally adapted genetic resources developed in harsh environments of the Brazilian Northwest (semi-arid) and maintained important traits for this region, such as parasite resistance, heat tolerance and high pelt quality. Genotypes (50K SNP chip) from seven Brazilian sheep breeds (5 hair and 2 coarse wool types) and 87 worldwide breeds were used to verify population structure, admixture and genetic diversity, using PCA and ADMIXTURE analyses. We constructed a phylogenetic tree and evaluated migration events between genetic groups using TREEMIX software. Brazilian Somali, a fat-tailed breed, was the unique breed with high relationship with East African breeds and formed a distinct cluster from other Brazilian breeds. This breed seems to contribute to formation of Santa Inês, Morada Nova and Brazilian Fat-tail breeds. Brazilian Blackbelly had a clear relationship with Barbados Blackbelly, which appeared as another group. Other Brazilian breeds seem to form a further genetic group with some recent admixtures. Morada Nova remained as a separate group, not showing a strong relationship with European or African breeds, only revealing a migration event from Sidaoun, an Algerian hair breed. Brazilian Fat-tail and Morada Nova share a common ancestor, but the first received introgressions from Brazilian Somali and Afrikaner breeds, explaining the fat-tail phenotype. Santa Inês received strong contribution from Bergamasca and had an admixed origin with recent introgressions from other breeds, mainly from Suffolk animals. In conclusion, Brazilian Somali and Brazilian Fat-tail are the most endangered sheep genetic resources in Brazil.

## Introduction

Cattle, sheep, goats and pigs were domesticated in the region from Central Anatolia to the north of the Zagros Mountains (southwest Asia), known as the Fertile Crescent (1). Sheep domestication started 11,000 years before the present (BP) (1) and played an important role in human society, spreading almost globally, following human migrations (2). Phylogeography of the genus *Ovis* is complex, involving several species and hybrids (3). Present day sheep (*Ovis orientalis aries*) are the product of two maternally distinct ancestral *Ovis gmelinii* populations and may be domesticated from Asiatic mouflon (*O. orientalis*), involving multiple independent domestication events in different geographic locations (1), mainly in the Middle East and North China. Strong historical human mediated gene flow across Eurasia has been observed (2). In addition, crossbreeding between wild and domestic populations has persisted and contributed to the high genetic diversity and admixture observed up to the present in sheep populations (4).

According to Lv et al. (2), the main migratory routes of sheep from Middle East to Eurasia and Africa included the Mediterranean, Danubian, Northern Europe and ancient sea trade routes to the Indian subcontinent, as well as routes of introduction and spread of sheep pastoralism in Africa. Thereafter, sheep populations were submitted to widely variable pastoral environments, which may have maintained natural selection pressure on sheep populations (2). Moreover, sheep were first reared for meat production, and later, for skin and milk production and, more recently, for wool (5), which contributes to different breeding goals and retained genetic diversity. Consequently, there are a broad spectrum of modern breeds adapted to a diverse range of environments and exhibiting specialized production of meat, milk, and fine wool (4).

Bakewell introduced the first idea of “breed” in England in the late 18th century and later the first herd book was opened in the mid 19^th^ century for Hereford cattle. Therefore, before this period, a “pure” breed vision or breed conservation efforts did not exist. In the 12^th^ century, Spain developed the Merino sheep, maintaining a monopoly on this breed with a very strong restriction to exportation of fine-wool animals (6). Probably, the settlers (mainly Portuguese and Spanish) brought animals with lower quality wool to the Americas (New World). Other animals may have come from Africa with the slave trade. There are no specific registers of sheep being taken to Brazil from Europe (7), but there is some reports of sheep introduced to Brazil through Paraguay and Argentina (6). Although controversial, the breeds supposedly brought to the Americas were Churra, Churra Bordaleira, Merino and Lacha (8).

Sheep in the Americas underwent natural selection and genetic drift during almost five centuries in certain environments. Brazilian hair breeds (Santa Inês, Morada Nova, Brazilian Somali, Brazilian Blackbelly, Cariri and Brazilian Fat-tail) initially were reared in Northeast region (hot and dry climate) in an extensive system with minimal care. In general, hair sheep breeds in Brazil have an important role in mutton production and their numbers are stable in recent years in comparison with wool breeds (9). Hair breeds also are known to produce the best skins in ruminants, which have a high demand in the clothing industry (10). These exotic but local adapted genetic resources have been jeopardized by the introduction of specialized breeds, often considered more productive and profitable. Therefore, the characterization of the genetic structure of these breeds is imperative for the preservation of these valuables genetic resources and developing viable conservation strategies.

Our study attempts to expand initial efforts (11,12) to understand the main origin and population structure of Brazilian local adapted sheep breeds. We were able to place Brazilian hair sheep breeds in a worldwide context using genomic information and better comprehend the relationship between these breeds.

## Materials and methods

### Genotypic data and Quality Control

Our initial data had genotypes of 4,014 animals, of which 1,149 animals were sampled in Brazil, 46 animals from Algeria (13) and 2819 animals from Sheep HapMap project - ISGC (International Sheep Genomics Consortium) (4). Brazilian samples came from Animal Germplasm Bank of Embrapa Genetic Resources and Biotechnology Centre. The Algerian breeds were included to test gene flow between Africa and South America, since some relationship was observed by Gaouar et al. (13).

All genotypes were obtained with the OvineSNP50 BeadChip (Illumina, San Diego, CA). Data was filtered using the following parameters in order: call rate per marker (<95%), minor allele frequency (MAF) <0.01 and sample call rate < 90%. Quality controls were performed using Golden Helix SNP & Variation Suite (SVS^®^) v. 8.6.0 (Golden Helix Inc., Bozeman, Montana, USA). In order to avoid bias in posterior analyses, we limited the number of animals per breed to a minimum of 5 and maximum of 50, and checked for genomic relatedness lower than 0.25. For breeds with more than 50 animals, samples were randomly chosen. Sample location diversity was maintained for Brazilian breeds. To reduce bias from linkage disequilibrium, LD pruning was used as follows: window size of 50 SNPs with window increment equal to 5 SNPs, r² > 0.5 with CHM method. The final data set had 2,549 animals (with 25,429 SNPs) from 94 breeds (S1 Table).

### Data analysis

Genotype Principal Component Analysis (PCA) was first conducted as a filtering step, identifying the breeds close to Brazilian breeds. The parameters were set to compute the first 10 components, normalizing each marker data by their actual standard deviation, utilizing an additive model and outlier removal up to 5 times, which was considered as more than 6 standard deviations, from 5 components.

Forty-four breeds were removed from the analysis based on the first and second component (S1 Fig). The data was reduced to 50 breeds and 1,202 animals. The proportion of variance explained by the third component of this PCA was close to the following components (S2 Fig) and a new PCA analysis was carried out. At this time, 22 breeds were removed (S3 Fig) and final data set was created with remaining 28 breeds and 490 animals (Table 1). Then, a third PCA analysis was realized to verify the relation of these breeds.

**Table 1.** Description of the 28 breeds used and results of inbreeding (F_IS_ statistics), standard deviation (SD) of F_IS_, observed (Ho) and expected (He) heterozygosity.

| Abb. | Breed | Country | Region | Classification | Tail | Tail lenght | Purpose | n | $F_{IS}$ | SD | $H_o$ |
| --- | --- | --- | --- | --- | --- | --- | --- | --- | --- | --- | --- |
| DOR | African Dorper | South_Africa | Africa | Hair | Fat | Short | Meat | 21 | 0.102 | 0.040 | 0.343 |
| AWD | African White Dorper | South_Africa | Africa | Hair | Fat | Short | Meat | 6 | 0.138 | 0.034 | 0.329 |
| BBB | Barbados BlackBelly | Caribbean | USA_Caribbean | Hair | Thin | Long | Meat | 24 | 0.164 | 0.091 | 0.319 |
| BAR | Barbarine Alg | Algeria | Africa | Coarse Wool | Fat | Long | Meat | 5 | 0.140 | 0.147 | 0.328 |
| BER | Berber Alg | Algeria | Africa | Coarse Wool | Thin | Long | Meat | 6 | 0.029 | 0.046 | 0.370 |
| BBA | Brazilian Bergamasca | Brazil | South_America | Coarse Wool | Thin | Long | Dual | 16 | 0.134 | 0.066 | 0.330 |
| BBN | Brazilian BlackBelly | Brazil | South_America | Hair | Thin | Long | Meat | 23 | 0.213 | 0.078 | 0.300 |
| BFT | Brazilian Fat-tail | Brazil | South_America | Hair | Fat | Long | Meat | 16 | 0.148 | 0.069 | 0.325 |
| BMN | Brazilian Morada Nova | Brazil | South_America | Hair | Thin | Long | Meat | 50 | 0.224 | 0.063 | 0.296 |
| BPT | Brazilian Pantaneira | Brazil | South_America | Coarse Wool | Thin | Long | Meat | 7 | 0.125 | 0.156 | 0.334 |
| BSI | Brazilian Santa Ines | Brazil | South_America | Hair | Thin | Long | Meat | 50 | 0.084 | 0.043 | 0.350 |
| BSO | Brazilian Somali | Brazil | South_America | Hair | Fat | Short | Meat | 25 | 0.227 | 0.105 | 0.295 |
| CHI | Chios | Spain | Europe | Coarse Wool | Fat | Long | Milk | 23 | 0.144 | 0.041 | 0.327 |
| COM | Comisana | Italy | Europe | Coarse Wool | Thin | Long | Milk | 24 | 0.023 | 0.014 | 0.373 |
| DMA | Dmen Alg | Algeria | Africa | Hair | Thin | Long | Meat | 5 | 0.055 | 0.042 | 0.361 |
| EMZ | Ethiopian Menz | Kenya | Africa | ? | Fat | Long | Dual | 34 | 0.156 | 0.037 | 0.322 |
| HAM | Hamra Alg | Algeria | Africa | Coarse Wool | Thin | Long | Meat | 6 | 0.072 | 0.031 | 0.354 |
| MCM | Macarthur Merino | Australia | Oceania | Fine Wool | Thin | Long | Wool | 10 | 0.382 | 0.037 | 0.236 |
| NA | Namaqua Afrikaner | South_Africa | Africa | Hair | Fat | Long | Meat | 12 | 0.241 | 0.026 | 0.290 |
| ODA | O.Djellal Alg | Algeria | Africa | Coarse Wool | Thin | Long | Meat | 6 | 0.014 | 0.010 | 0.376 |
| RM | Red Maasai | Kenya | Africa | Hair | Fat | ? | Meat | 45 | 0.145 | 0.031 | 0.326 |
| REM | Rembi Alg | Algeria | Africa | Coarse Wool | Thin | Long | Meat | 6 | 0.002 | 0.009 | 0.381 |
| RA | Ronderib Afrikaner | South_Africa | Africa | Hair | Fat | Long | Meat | 17 | 0.174 | 0.068 | 0.315 |
| SAK | Sakiz | Turkey | Middle_East | Coarse Wool | Thin | Long | Wool | 22 | 0.123 | 0.053 | 0.335 |
| SID | Sidaoun Alg | Algeria | Africa | Hair | Thin | Long | Meat | 6 | 0.127 | 0.062 | 0.333 |
| STE | St Elizabeth | Caribbean | USA_Caribbean | ? | Thin | ? | ? | 10 | 0.018 | 0.032 | 0.375 |
| SUF | Suffolk | England | Europe | Medium Wool | Thin | Short | Meat | 9 | 0.076 | 0.063 | 0.352 |
| TAZ | Tazgezawth Alg | Algeria | Africa | Coarse Wool | Thin | Long | Meat | 6 | 0.182 | 0.122 | 0.312 |

Abb.: abbreviation of the breed name; n: number of animals evaluated in each breed; $F_{IS}$ : inbreeding statistics (Wright's F statistics); SD: standard deviation of $F_{IS}$ ; $H_o$ : observed heterozygosity. Expected heterozygosity was 0.38 for all breeds.

Genetic diversity inside populations was estimated from inbreeding coefficient (F_IS_), observed proportion of heterozygote genotypes per individual (H_o_) and expected heterozygosity (H_e_) using SNP & Variation Suite v8.7 (Golden Helix, Inc., Bozeman, MT, www.goldenhelix.com).

Pair-wise fixation index (F_ST_) was calculated as a first measure of genetic diversity between populations. Genetic relationships among breeds and level of admixture were evaluated through the model-based clustering algorithm implemented in the software Admixture v. 1.3.0 (14). The cross-validation procedure (10-fold) was carried out to estimate prediction errors for each K value (from 2 to 30). The value of K that minimizes this estimated prediction error represents the best predictive accuracy for the model. Individual coefficients of membership for each K cluster produced by Admixture were visualized using the on-line Clumpak server with the feature Distruct for many K’s (15).

To refine the genetic structure of Brazilian hair sheep, we performed a second cluster analysis with only 7 breeds with closest average F_ST_ values (Comisana, Brazilian Suffolk, Brazilian Santa Inês, Brazilian Morada Nova, Brazilian Bergamasca, Brazilian Pantaneira and Brazilian Fat-tail) using Admixture v. 1.3.0 (14). We used K from 2 to 9 and performed the same cross-validation, best-K identification and visualization procedures.

Treemix software, a composite likelihood maximization tree-based approach, was used to reconstruct historical relationships between the analyzed populations and to test for the presence of gene flow (16). The software was run on two datasets similar to Admixture (with the previous 28 breeds and only the 7 breeds). Soay breed was used as external group in both analyses, following the observed by Kijas et al. (4). Soay sheep inhabits the island of St. Kilda off the northwest Scotland (geographically isolated) and was previously identified as firmly linked to Mediterranean and Asiatic Mouflon, considered ancestors of domestic sheep (5).

A variable number of migration events (M) (0, 1, 2, 3, 4, 5, 10, 20, 30, 40 and 50) were tested. The value of M that had the highest log-likelihood indicated the most predictive model. In order to improve gene flow analysis, three-population (f_3_) and four-population (f_4_) tests (17) were carried out using Treemix software. Significance of results is evaluated by weighted block jackknife to obtain a mean Z-score. The f_3_ tests are presented as f_3_(A; B, C), where a significantly negative value of the f_3_ statistic implies that population A is admixed from populations B and C. The f_4_ statistics are in the form f_4_(A, B; C, D) where: a zero value indicates that there is more gene flow between A and B and between C and D than the other possible combinations in the tree; a negative value means that A and D, B and C show the closest relationship; and a positive value means that A and C, B and D are the closest relationship in the tree. Both tests were carried out using blocks of 500 SNPs and 100 SNPs. The results between them were similar (S2 to S4 Tables). A complete explanation about the calculation and evaluation of the f3 and f4 can be seen in Peter (18) and Reich et al. (17).

## Results

The first PCA analysis (S1 Fig) showed divergence between a group of European and Oceanian breeds from Asiatic breeds. As both groups were distant from Brazilian breeds, they were removed from the analysis. The second PCA (S3 Fig) demonstrates a separation of South American and African populations. In this analysis, a group of breeds mainly from Europe, Middle East and wool breeds of South America remained close to zero, and they were removed from further analyses.

The final PCA with 28 breeds (Table 1) demonstrated a relationship between Brazilian Somali and some African breeds (Fig 1 and S4 Fig). The Morada Nova breed was seen partially isolated in both components. Brazilian breeds (Santa Inês, Pantaneira, Bergamasca, Blackbelly and Suffolk), Comisana, Barbados Blackbelly and St. Elizabeth were seen relatively close.

**Fig 1.**
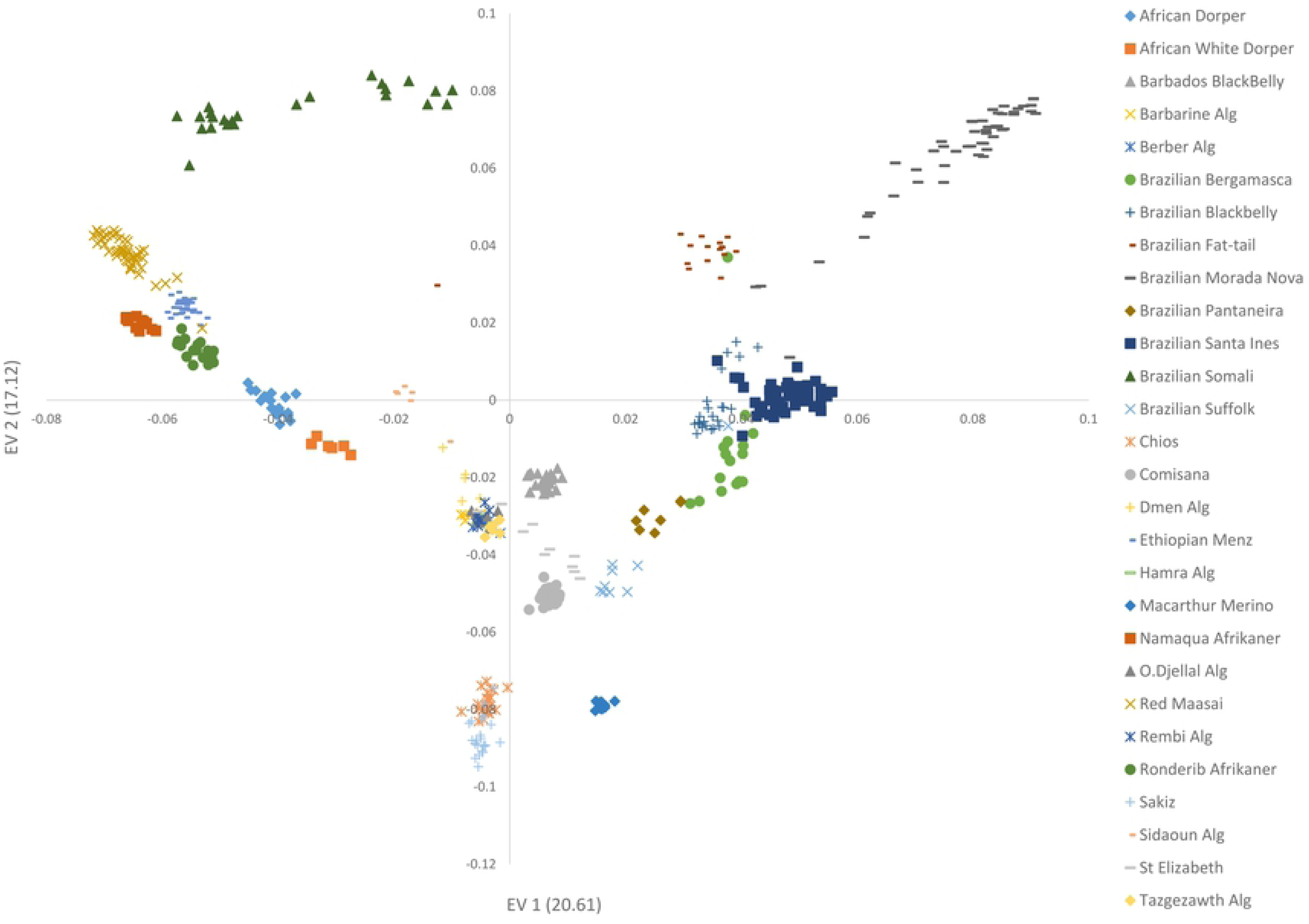
Principal component analysis (PCA) using genomic data of 28 breeds close to Brazilian hair sheep breeds. The number between parentheses in each axes means the amount of the variance explained by each eigenvector (EV).

Within breed genetic diversity for the 28 breeds is shown in Table 1 (Ho, He and Fis). In general, Brazilian hair breeds showed high Fis (inbreeding coefficient), except for Santa Inês. This can be related to the F_ST_ results, where the Brazilian breeds generally presented higher values than the others.

The Brazilian Blackbelly (BBN) showed lowest F_ST_ (Table 2) with Barbados BlackBelly (BBB) (0.116) and Santa Inês (BSI) (0.119). The BBN and BBB are phenotypically very similar, so this proximity was expected. Brazilian Fat-tail (BFT) showed some proximity with Algerian breeds (Berber, O.Djellal and Rembi), in particular with fat-tailed Barbarine (13).

**Table 2.**
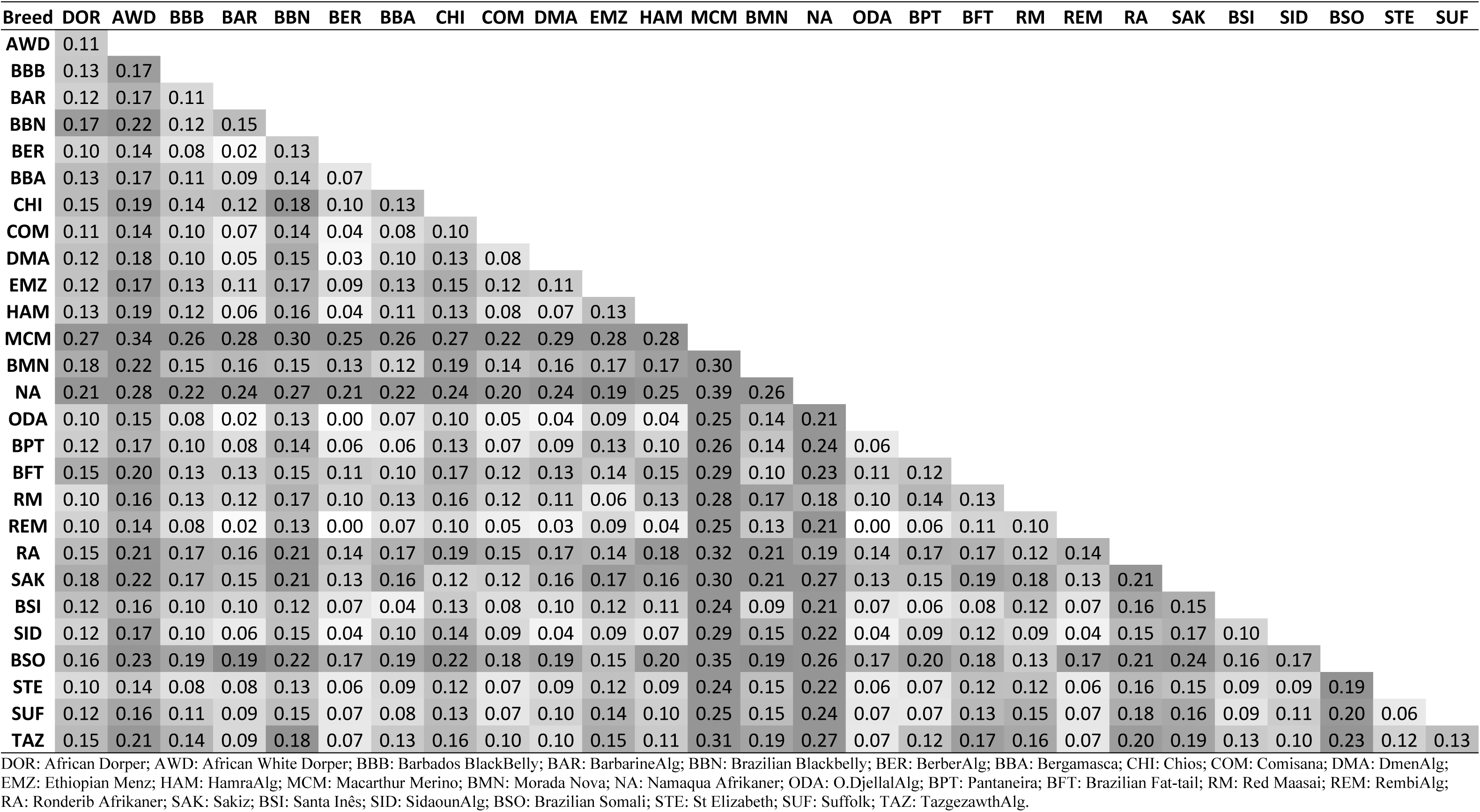
Pair-wise fixation index (F_ST_) results for pairs of the 28 sheep breeds.

| Breed | DOR | AWD | BBB | BAR | BBN | BER | BBA | CHI | COM | DMA | EMZ | HAM | MCM | BMN | NA | ODA | BPT | BFT | RM | REM | RA | SAK | BSI | SID | BSO | STE | SUF |
| --- | --- | --- | --- | --- | --- | --- | --- | --- | --- | --- | --- | --- | --- | --- | --- | --- | --- | --- | --- | --- | --- | --- | --- | --- | --- | --- | --- |
| AWD | 0.11 |  |  |  |  |  |  |  |  |  |  |  |  |  |  |  |  |  |  |  |  |  |  |  |  |  |  |
| BBB | 0.13 | 0.17 |  |  |  |  |  |  |  |  |  |  |  |  |  |  |  |  |  |  |  |  |  |  |  |  |  |
| BAR | 0.12 | 0.17 | 0.11 |  |  |  |  |  |  |  |  |  |  |  |  |  |  |  |  |  |  |  |  |  |  |  |  |
| BBN | 0.17 | 0.22 | 0.12 | 0.15 |  |  |  |  |  |  |  |  |  |  |  |  |  |  |  |  |  |  |  |  |  |  |  |
| BER | 0.10 | 0.14 | 0.08 | 0.02 | 0.13 |  |  |  |  |  |  |  |  |  |  |  |  |  |  |  |  |  |  |  |  |  |  |
| BBA | 0.13 | 0.17 | 0.11 | 0.09 | 0.14 | 0.07 |  |  |  |  |  |  |  |  |  |  |  |  |  |  |  |  |  |  |  |  |  |
| CHI | 0.15 | 0.19 | 0.14 | 0.12 | 0.18 | 0.10 | 0.13 |  |  |  |  |  |  |  |  |  |  |  |  |  |  |  |  |  |  |  |  |
| COM | 0.11 | 0.14 | 0.10 | 0.07 | 0.14 | 0.04 | 0.08 | 0.10 |  |  |  |  |  |  |  |  |  |  |  |  |  |  |  |  |  |  |  |
| DMA | 0.12 | 0.18 | 0.10 | 0.05 | 0.15 | 0.03 | 0.10 | 0.13 | 0.08 |  |  |  |  |  |  |  |  |  |  |  |  |  |  |  |  |  |  |
| EMZ | 0.12 | 0.17 | 0.13 | 0.11 | 0.17 | 0.09 | 0.13 | 0.15 | 0.12 | 0.11 |  |  |  |  |  |  |  |  |  |  |  |  |  |  |  |  |  |
| HAM | 0.13 | 0.19 | 0.12 | 0.06 | 0.16 | 0.04 | 0.11 | 0.13 | 0.08 | 0.07 | 0.13 |  |  |  |  |  |  |  |  |  |  |  |  |  |  |  |  |
| MCM | 0.27 | 0.34 | 0.26 | 0.28 | 0.30 | 0.25 | 0.26 | 0.27 | 0.22 | 0.29 | 0.28 | 0.28 |  |  |  |  |  |  |  |  |  |  |  |  |  |  |  |
| BMN | 0.18 | 0.22 | 0.15 | 0.16 | 0.15 | 0.13 | 0.12 | 0.19 | 0.14 | 0.16 | 0.17 | 0.17 | 0.30 |  |  |  |  |  |  |  |  |  |  |  |  |  |  |
| NA | 0.21 | 0.28 | 0.22 | 0.24 | 0.27 | 0.21 | 0.22 | 0.24 | 0.20 | 0.24 | 0.19 | 0.25 | 0.39 | 0.26 |  |  |  |  |  |  |  |  |  |  |  |  |  |
| ODA | 0.10 | 0.15 | 0.08 | 0.02 | 0.13 | 0.00 | 0.07 | 0.10 | 0.05 | 0.04 | 0.09 | 0.04 | 0.25 | 0.14 | 0.21 |  |  |  |  |  |  |  |  |  |  |  |  |
| BPT | 0.12 | 0.17 | 0.10 | 0.08 | 0.14 | 0.06 | 0.06 | 0.13 | 0.07 | 0.09 | 0.13 | 0.10 | 0.26 | 0.14 | 0.24 | 0.06 |  |  |  |  |  |  |  |  |  |  |  |
| BFT | 0.15 | 0.20 | 0.13 | 0.13 | 0.15 | 0.11 | 0.10 | 0.17 | 0.12 | 0.13 | 0.14 | 0.15 | 0.29 | 0.10 | 0.23 | 0.11 | 0.12 |  |  |  |  |  |  |  |  |  |  |
| RM | 0.10 | 0.16 | 0.13 | 0.12 | 0.17 | 0.10 | 0.13 | 0.16 | 0.12 | 0.11 | 0.06 | 0.13 | 0.28 | 0.17 | 0.18 | 0.10 | 0.14 | 0.13 |  |  |  |  |  |  |  |  |  |
| REM | 0.10 | 0.14 | 0.08 | 0.02 | 0.13 | 0.00 | 0.07 | 0.10 | 0.05 | 0.03 | 0.09 | 0.04 | 0.25 | 0.13 | 0.21 | 0.00 | 0.06 | 0.11 | 0.10 |  |  |  |  |  |  |  |  |
| RA | 0.15 | 0.21 | 0.17 | 0.16 | 0.21 | 0.14 | 0.17 | 0.19 | 0.15 | 0.17 | 0.14 | 0.18 | 0.32 | 0.21 | 0.19 | 0.14 | 0.17 | 0.17 | 0.12 | 0.14 |  |  |  |  |  |  |  |
| SAK | 0.18 | 0.22 | 0.17 | 0.15 | 0.21 | 0.13 | 0.16 | 0.12 | 0.12 | 0.16 | 0.17 | 0.16 | 0.30 | 0.21 | 0.27 | 0.13 | 0.15 | 0.19 | 0.18 | 0.13 | 0.21 |  |  |  |  |  |  |
| BSI | 0.12 | 0.16 | 0.10 | 0.10 | 0.12 | 0.07 | 0.04 | 0.13 | 0.08 | 0.10 | 0.12 | 0.11 | 0.24 | 0.09 | 0.21 | 0.07 | 0.06 | 0.08 | 0.12 | 0.07 | 0.16 | 0.15 |  |  |  |  |  |
| SID | 0.12 | 0.17 | 0.10 | 0.06 | 0.15 | 0.04 | 0.10 | 0.14 | 0.09 | 0.04 | 0.09 | 0.07 | 0.29 | 0.15 | 0.22 | 0.04 | 0.09 | 0.12 | 0.09 | 0.04 | 0.15 | 0.17 | 0.10 |  |  |  |  |
| BSO | 0.16 | 0.23 | 0.19 | 0.19 | 0.22 | 0.17 | 0.19 | 0.22 | 0.18 | 0.19 | 0.15 | 0.20 | 0.35 | 0.19 | 0.26 | 0.17 | 0.20 | 0.18 | 0.13 | 0.17 | 0.21 | 0.24 | 0.16 | 0.17 |  |  |  |
| STE | 0.10 | 0.14 | 0.08 | 0.08 | 0.13 | 0.06 | 0.09 | 0.12 | 0.07 | 0.09 | 0.12 | 0.09 | 0.24 | 0.15 | 0.22 | 0.06 | 0.07 | 0.12 | 0.12 | 0.06 | 0.16 | 0.15 | 0.09 | 0.09 | 0.19 |  |  |
| SUF | 0.12 | 0.16 | 0.11 | 0.09 | 0.15 | 0.07 | 0.08 | 0.13 | 0.07 | 0.10 | 0.14 | 0.10 | 0.25 | 0.15 | 0.24 | 0.07 | 0.07 | 0.13 | 0.15 | 0.07 | 0.18 | 0.16 | 0.09 | 0.11 | 0.20 | 0.06 |  |
| TAZ | 0.15 | 0.21 | 0.14 | 0.09 | 0.18 | 0.07 | 0.13 | 0.16 | 0.10 | 0.10 | 0.15 | 0.11 | 0.31 | 0.19 | 0.27 | 0.07 | 0.12 | 0.17 | 0.16 | 0.07 | 0.20 | 0.19 | 0.13 | 0.10 | 0.23 | 0.12 | 0.13 |

DOR: African Dorper; AWD: African White Dorper; BBB: Barbados BlackBelly; BAR: BarbarineAlg; BBN: Brazilian Blackbelly; BER: BerberAlg; BBA: Bergamasca; CHI: Chios; COM: Comisana; DMA: DmenAlg; EMZ: Ethiopian Menz; HAM: HamraAlg; MCM: Macarthur Merino; BMN: Morada Nova; NA: Namaqua Afrikaner; ODA: O.DjellalAlg; BPT: Pantaneira; BFT: Brazilian Fat-tail; RM: Red Maasai; REM: RembiAlg; RA: Ronderib Afrikaner; SAK: Sakiz; BSI: Santa Inês; SID: SidaounAlg; BSO: Brazilian Somali; STE: St Elizabeth; SUF: Suffolk; TAZ: TazgezawthAlg.

Brazilian Morada Nova (BMN), in general, showed high F_ST_ with other breeds and only showed a close relationship with BSI, Bergamasca and BFT. Brazilian Somali showed high FST with all breeds, which can be explained by the high inbreeding (FIS) of this group (0.23). Brazilian Bergamasca (BBA) showed low F_ST_ (<0.10) with Algerian breeds (Barbarine, Berber, Dmen, O. Djellal and Rembi). BBA also showed low F_ST_ with Comisana, St Elizabeth and Suffolk. Nevertheless, the Bergamasca had the lowest F_ST_ with Pantaneira (BPT) and BSI. Pantaneira (BPT), which is a coarse wool breed reared in a temporary flooded ecosystem of Brazil, called Pantanal, seems to be very close to Bergamasca. Santa Inês (BSI) showed low F_ST_ (<0.10) with Algerian, Comisana and Suffolk breeds. Nevertheless, BSI had low F_ST_ with other Brazilian breeds and the lowest with BBA.

The results of ADMIXTURE with 28 breeds showed, at K=3, African, European and Brazilian components in genetic structure of these populations (Fig 2). The best K was considered as 16, while from 16 to 18 the prediction error was very close (S5 Fig). At K=16, Brazilian Somali, Brazilian Fat-tail, Brazilian Blackbelly and Morada Nova differentiated from other groups and between themselves. Santa Inês showed some admixture with Bergamasca, Pantaneira, Brazilian Fat-tail, Comisana and Suffolk. Brazilian Blackbelly demonstrates clearly two groups of animals in this population. At higher K’s (S6 Fig), Morada Nova and Santa Inês show substructure that reflect geographic and varietal differences.

**Fig 2.**
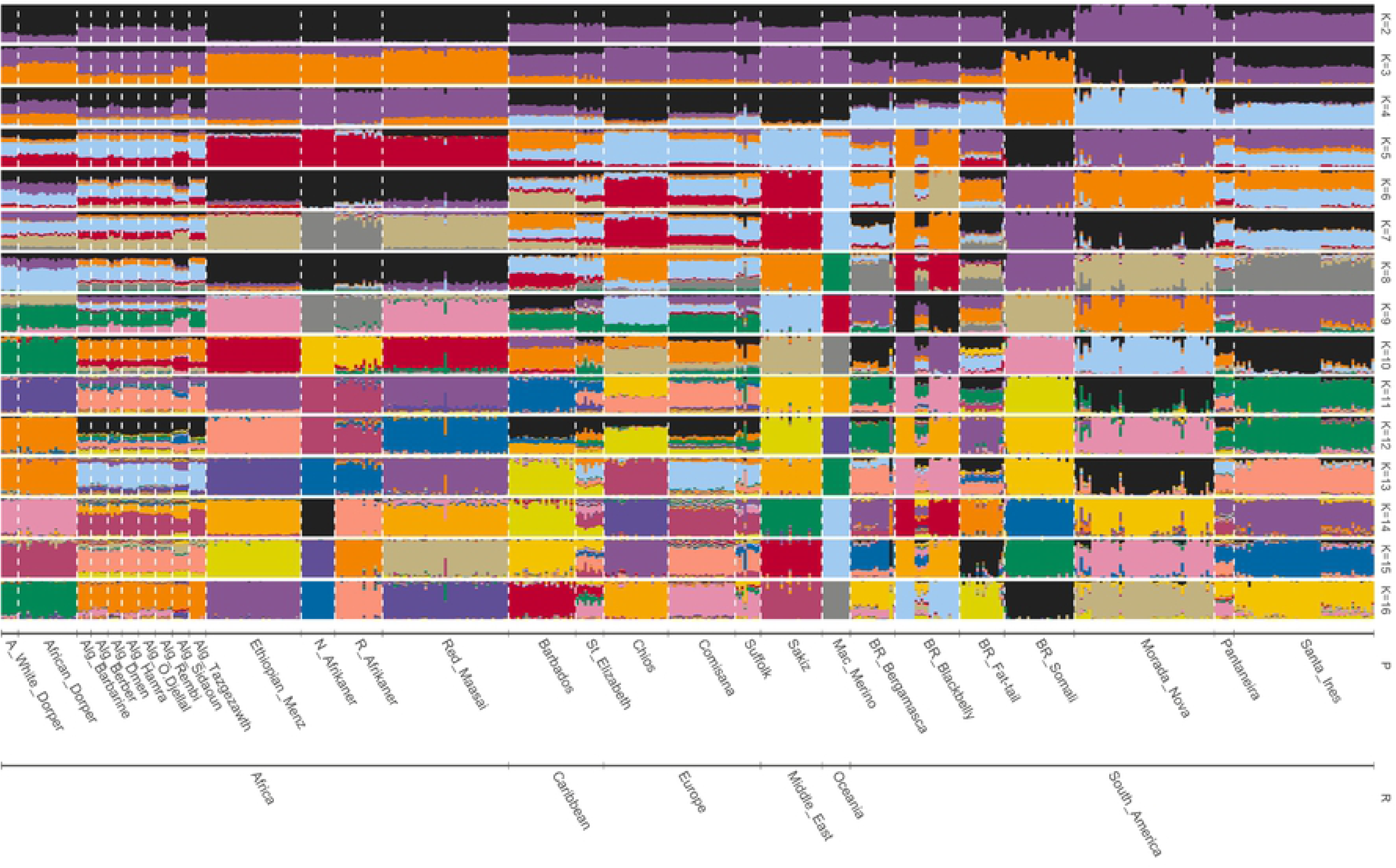
Clustering (ADMIXTURE software) based on genotypic data from 28 breeds close to Brazilian hair sheep breeds. (K) number of clusters, from 2 to 16. Populations (P) grouped according to region (R).

The best K of admixture analysis with 7 breeds was equal to 4 (S7 Fig). At K=4, the plot shows that Morada Nova and Brazilian Fat-tail are in separate groups (Fig 3). In addition, Comisana differentiated from the others. The genetic structure of Santa Inês seems be composed mainly by Bergamasca, with minor contribution of Morada Nova, Brazilian Fat-tail and Suffolk in some animals. Again, a subdivision inside the breed was observed.

**Fig 3.**
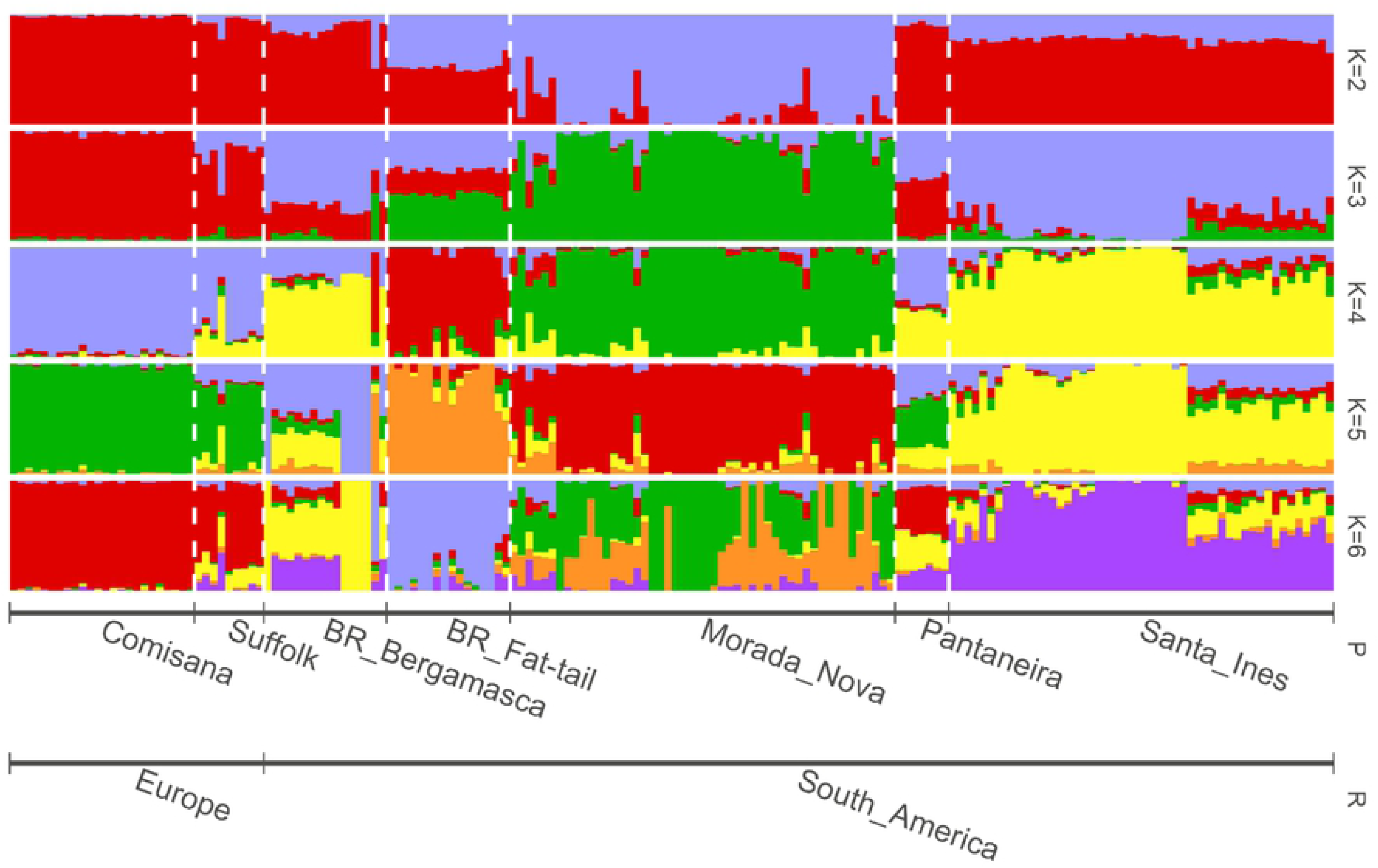
Model-based clustering (ADMIXTURE) using genotypic data from 7 sheep breeds with close relationship. (K) number of clusters, from 2 to 6. The best prediction model was K = 4. Populations (P) grouped according to region (R).

The inferred trees with 29 breeds in TREEMIX software (Fig 4 and S8 Fig) showed Brazilian Somali in an African branch, next to Red Maasai and Ethiopian Menz. Brazilian Blackbelly remained close to the Barbados BlackBelly as observed in other analyses. Remaining Brazilian and Algerian breeds formed two separate branches.

**Fig 4.**
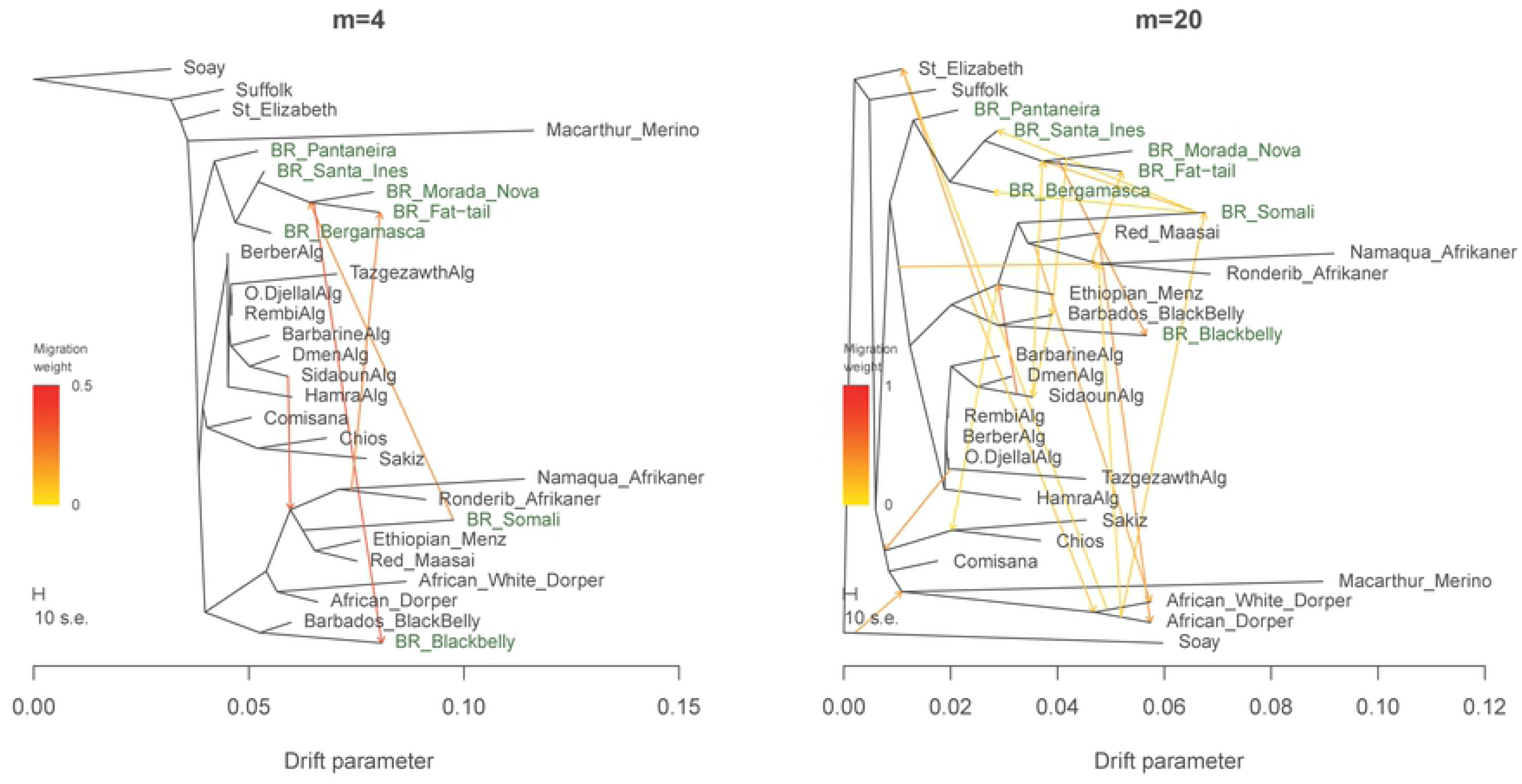
Inferred trees from 29 sheep breeds using TREEMIX software. The trees were construct with 4 and 20 migration events (m = 4 and m = 20, respectively). Soay breed was used as root. Brazilian breeds highlighted in green.

The evaluation of the number of migration events showed an increasing ln(likelihood), however the improvement after 20 migration events was small (S9 Fig). As the number of migrations increased, it becomes more difficult to understand the relationships. According to Pickrell and Pritchard (16), sometimes it may be preferable to stop adding migration events well before the maximum likelihood point so that the resulting graph remains interpretable. Therefore, we show the inferred tree with 20 migration events due the small increment in likelihood beyond this point.

With four migration events, a contribution of an ancestral Ronderib and Namaqua Afrikaner to Brazilian Fat-tail constitution is highlighted and some migration events between Brazilian breeds, such as Brazilian Somali to an ancestor of Morada Nova and Brazilian Fat-tail, and from this ancestor to Brazilian Blackbelly. The tree with 20 migration events showed main migration events involving Brazilian breeds were: Brazilian Somali to BSI, BMN, BFT and BBA; Afrikaner breeds to BFT; Sidaoun (Algerian breed) to BMN and BFT ancestral; BMN to BBN (the highest weight); as well as African Dorper to the actual Brazilian Somali, which can be related to recent introgression.

Observing the tree residuals involving Brazilian breeds, the tree with 20 migrations underestimates the observed covariance between Brazilian Somali and Pantaneira, as well as Suffolk with Morada Nova and Pantaneira (S10 Fig). The tree using the same 7 breeds analyzed in ADMIXTURE (Fig 5) had the best model with 3 migration events (S11 Fig). The migrations defined a relationship between Santa Ines and Morada Nova, as well as a migration from Comisana branch to an ancestral of BFT and BMN. The residuals (S12 Fig) shows that the tree underestimates the relationship between Pantaneira and Bergamasca, and slightly overestimates the covariance between Suffolk and Bergamasca.

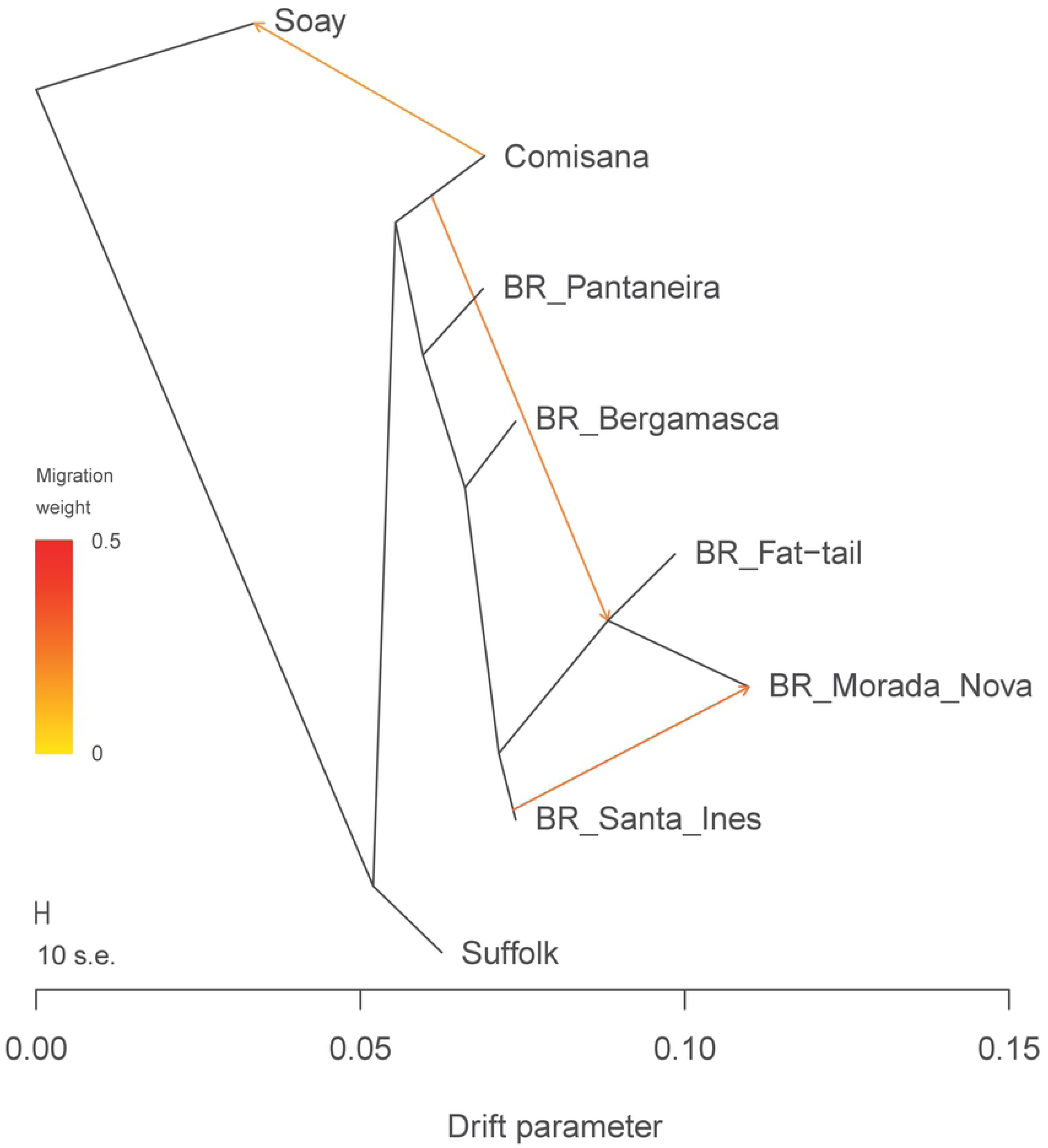

The results of f_3_ statistics with the 28 breeds (S2 Table) showed the great admixture process between the Algerian breeds, with the Sidaoun breed contributing to genetic composition of Berber, O. Djellal and Rembi. The f_3_ (Santa Ines; Morada Nova, Bergamasca) was the lowest values when analyzing only the 7 breeds (S4 Table), but remained positive. This value expresses the length of the branch in the unrooted three-populations phylogeny leading to A from the internal node (18). Therefore, this result supports the hypothesis of the Santa Ines breed coming from crossbreeding between Morada Nova and Bergamasca.

The f_4_ statistic (S3 and S5 Tables) showed greater relationship between Comisana and Suffolk when compared to relationship between Santa Inês, Morada Nova, Pantaneira, Brazilian Fat-tail and Bergamasca. Comisana and Bergamasca rarely grouped together in the f_4_ statistics, despite being two Italian breeds. Observing the f_4_ statistic with Soay (considered as outgroup), Comisana grouped with Soay, and Suffolk showed higher relationship with Pantaneira, Bergamasca, Morada Nova and Santa Inês (S3 Table). Therefore, Suffolk animals had a higher relationship with Brazilian breeds than with Comisana, as the results of PCA (Fig. 1) and ADMIXTURE (Fig. 2 and 3) suggested. Between Brazilian breeds, Pantaneira seems to be the breed with the highest relationship with Suffolk and Comisana.

## Discussion

Brazilian hair sheep breeds have a mixed origin. Brazilian Somali has a strong East African origin. Brazilian Blackbelly had an expected relationship with Barbados BlackBelly and they are more related with East African than European breeds. The remaining Brazilian hair sheep breeds seem to form a unique and differentiated group that has a suggested mixed origin from Mediterranean and West African regions.

### Origin of Brazilian hair sheep

During the Colonial period, the Portuguese had strong commercial ties with Indian colony. Long sea journeys required a good stock of food and sheep were a likely source. Afterwards, goats such as Bhuj and Jamnapari arrived in Brazil with Zebu cattle importers in the early 20th century (26). Other importers may have also brought sheep which were not registered due to low commercial value and perceived lack of importance (27). In the first PCA (S1 Fig), some Asian breeds (Indian Garole, Bangladeshi Garole, Sumatra, Deccani, Tibetan, Changthangi and Garut) were evaluated. However, these breeds were removed in the first PCA analysis, showing high genetic distance from Brazilian hair breeds.

In the reduced data set with 28 breeds, genetic divisions were detected separating European, African and Brazilian sheep (K=3, Fig 4). Kijas et al. (4) also found sheep from the Americas (Brazil and Caribbean) cluster separately from European, African, or Asian populations. These authors stated a genetic origin for Caribbean breeds in common with African animals mixed with those of Mediterranean Europe. Surprisingly, the only Brazilian breed in our study with strong African origin is the Brazilian Somali. The other Brazilian breeds seem to be formed through migration processes and are related with ancestral formation of Mediterranean and African breeds, which is expected due the Colonial process of Portugal and Spain followed by slave migration from Africa.

Brazilian Blackbelly animals have a brown body coat colour and black coat in the belly and internal part of legs, being very similar to the Barbados Blackbelly, therefore the relationship between them was expected. It is reported that this breed was imported from West Africa with slaves and were later crossed with imported wool sheep from Europe, selecting against the presence of wool (19). Between 1630 and 1654, during the Dutch domination of the Brazilian Northeast, there was a genetic group of sheep called Jaguaribe which was similar to African breeds and probably is an ancestral of some Brazilian hair breeds (20). Dutch settlers were driven out from Brazil and went to Antilles islands, carrying sugar cane production and, probably, the ancestral hair sheep that originated Barbados Blackbelly. The tree plot (Fig 6) supports this hypothesis showing the Brazilian Blackbelly and Barbados Blackbelly branch. Spangler et al. (21) also enforces the link between West African (Djallonké sheep) and Caribbean sheep breeds.

Brazilian Somali and Brazilian Fat-tail are fat-tail hair breeds and seem to have different origins. The Brazilian Somali was close to East African breeds in all analyses, while Brazilian Fat-tail remained close to other Brazilian breeds. Both breeds represent unique genetic architecture in cluster analysis, forming an exclusive cluster from K=4 for Brazilian Somali and K=14 for Brazilian Fat-tail. These results may be related to the small number of animals remaining in each breed (22,23) and, consequently, the strong genetic drift suffered in both cases.

Brazilian Somali were related to a common ancestor of the Red Maasai, Ethiopian Menz and Afrikaner breeds (Ronderib and Namaqua), which is also a fat-tailed group. Their black head and white body are similar to some African breeds such as Dorper, BlackHead Persian and others. The Brazilian Somali probably has its origin in the horn of Africa and is thought to have the Urial as its ancestor (23). This animal first arrived in Brazil in 1939, brought by farmers from Rio de Janeiro State, but the breed thrive in the drier and hotter climates found in the northeast of the country (24).

Brazilian Fat-tail shares a common ancestor with Morada Nova (Fig 6), which is expected as both are reared in similar regions (22). Brazilian Fat-tail probably resulted from crossbreeding Brazilian Somali and Afrikaner breeds (migration events), which are also fat-tailed breeds.

In the cluster analysis (Fig 4 and 5), the Morada Nova breed appears in a specific cluster. Morada Nova branch received a migration event from Sidaoun, an Algerian breed (m=20, Fig 6). This breed, also known as Targui (Targuia, Sidaou or Sidaho) is a hair sheep exploited under nomadic conditions by the Tuareg people in the southern part of Algeria (Central Sahara) (25). Originated from Mali, it is a highly rustic breed, well adapted to walk long distances “transhumance” and harsh climatic conditions (25). A relationship between Morada Nova and a Nigerian breed (Djallonké or Dwarf) was also seen (21), disclosing the possible west African origin of Morada Nova. Lima Pereira (37) states categorically that the larger variety of sheep in Angola is the ancestor of the Morada Nova; as they both lack the mane of the Fouta Djallon.

Santa Inês is the main commercial hair breed in Brazil; however, its origin is still unknown. ARCO (Brazilian Association of Sheep breeding - www.arcoovinos.com.br) states as originated from crossing between Bergamasca, Morada Nova, Brazilian Somali and other undefined sheep. They also state that the wool remnants are due to the Bergamasca, while its lack of wool and coat types are due to the Morada Nova. According to ARCO, the participation of Brazilian Somali could be observed in the fat around the tail head when the animal is very fat. The present study showed a migration event from Brazilian Somali to Santa Inês (m=20; Fig 6), corroborating with this statement.

Sheep with similar traits to the Morada Nova and Santa Inês exist in several Western and Central African countries, perhaps in the cluster analysis of this study (Fig 4 and 5), these two breeds did not show any African component. At present, the main hypothesis is that these African animals would have been crossed with elite animals from Iberian breeds such as Churra and Bordaleira (28). The Churra breed is in the initial data of this study, but it was removed in the second PCA. Therefore, this breed did not show a close relationship to Santa Inês or Morada Nova as expected.

The Pelibuey breed (which means skin of cattle, “Pêlo-de-boi”) found in the USA, Mexico as well as other America countries is phenotypically similar to the Santa Inês (29,30). Its origin may also be related to animals that left the Brazilian Northeast by Dutch colonialism migration, similar to that seen with Brazilian Blackbelly and Barbados BlackBelly.

The Bergamasca breed, which has Italian origin, is clearly the main composition of the Santa Inês animals (Fig 5). Some previous studies also observed the proximity between these breeds (6,11,31,32). There is no specific records about the importation of this breed to South America, so little is known about their arrival (date, quantity of animals, etc) and rearing conditions in Brazil (where these animals were placed). Nevertheless, Bergamasca arrived in Brazil with the Italian settlers from the late 19th century up to the 1930s (33).

In a study with Italian breeds (35), two Alpine breeds, Engadine Red and Alpagota were the closest breeds to the Bergamasca/Biellese group. Bergamasca is known to have been used as a crossing breed in northern Italy, southern Germany, Austria, Slovenia and central Italy because of its large frame (35). Probably, because of the same reason, Bergamasca animals were crossed with Morada Nova ancestors (local ecotype), and them this local sheep was backcrossed and selected against the presence of wool and/or natural selection to the harsh environmental conditions of Brazilian Northwest, lacking the wool in the same way. Other possibility is that Bergamasca was crossed with local hair sheep to improve fitness traits and them they was backcrossed to maintain Bergamasca breed. It is necessary to evaluate the relationship between Brazilian and Italian Bergamasca to comprehend better the structure of this breed.

### Conservation impact

Paiva et al. (12) showed that locally adapted sheep breeds in Brazil were closely related, which may be due to crossbreeding in the past in association with genetic drift. These close relationships were also seen here in both admixture and tree plot. Only Brazilian Somali and Brazilian Blackbelly show some genetic divergence appearing in the African and Blackbelly branches respectively, but the migration analysis (both m=4 and m=20; Fig 6) demonstrated some crossbreeding of these breeds with other Brazilian breeds.

Brazilian Somali animals has been used extensively in absorbent crossbreeding to create the “Brazilian Dorper” (23). A migration event was observed from the Dorper branch to Brazilian Somali (Fig 6), supporting this recent introgression. As Brazilian farmers consider Dorper as more productive, Brazilian Somali is threatened by extinction. Ianella et al. (38), using 13 sheep breeds in Brazil, observed Brazilian Somali had a low level of susceptibility to Scrapie based on PRNP allele frequencies, while Dorper had the highest level. Therefore, the crossbreeding may proportionate a genetic erosion and predisposes the occurrence of diseases that have not previously occurred in Brazilian locally adapted sheep.

Brazilian Fat-tail are reared in the hottest climate with the lowest precipitation index and consequently with the highest temperature and humidity index between all sheep breeds reared in Brazil (22). This breed had the lowest number of flocks and the lowest distance from mid-point of breed occurrence (22). These previous findings with the unique genetic structure (K=4; Fig 5) observed in the present study highlights the demand for genetic conservation efforts for this breed.

The Morada Nova breed is reared in the Northwest region of Brazil, mainly in Ceará state (22). Animals have small size with small and short ears, the main coat colour is reddish brown, but some variation exists with a white lineage (11). Breeders use Morada Nova animals for meat production, but only a recent restructuring of the Morada Nova Breed Association in 2008 provided the basis for the establishment of a local community-based breeding program coordinated by Embrapa Sheep and Goat Research Centre (CNPCO). The actual herds remain next to the center of origin (22) and, as showed here, have a high inbreeding coefficient. Therefore, this genetic group established a unique genetic constitution probably due genetic drift and natural selection to the harsh environmental conditions in the region of origin.

Morada Nova had a high allelic diversity for the *FecG^E^* allele (GDF9 gene), which has been shown to increase litter size (40). This allele had previously been described only in Brazilian Santa Inês sheep (41). These studies suggest that this mutation might be associated only with Brazilian locally adapted sheep breeds. Therefore, these results highlight the relevance of applying conservation efforts for this genetic resource.

There are substantial differences between the Red and White varieties of Morada Nova, which should be used as separate genetic resources to improve conservation programs (39). This subgrouping inside the breed can also be seen here (K>25; S4 Fig). Therefore, these findings suggest the possibility of development of an animal breeding program with the use of these lineages inside the breed.

Santa Inês breed has gone expressive expansion in last decade with some crossbreeding with other populations (6,11,31) and nowadays is the main commercial hair breed in Brazil (22). Paiva et al. (12) stated the hypothesis of existence of the “old Santa Inês” and the “New Santa Inês”, due to recent crossbreeding. Old Santa Inês may be classified as smaller and more rustic animals which were predominant in the 80s and 90s (42). The New Santa Inês have a large body (main rump and legs) which appeared in a large portion of the population in only a few years, which is more likely due to crossbreeding than within breed selection, even considering that the breed did not have an official breeding program. The color of the breed also changed and is now predominantly brown or black whereas before various coat colors could be found (31). The risk associated with this “unknown” crossbreeding are the loss of important traits such as gastro-intestinal parasite resistance (43), pelt quality (10), heat resistance (44), and also the insertion of non-desirable traits such as scrapie susceptibility (38), as explained earlier for Brazilian Somali.

Some previous studies (11,31) stated a possible crossbreeding of Santa Inês with meat-selected breeds, mainly with Suffolk. This hypothesis can be confirmed here as the genetic structure of some animals has a contribution from Suffolk (Fig. 5), which was confirmed in the f_4_ statistics. In the present study, at higher K’s (S4 Fig), a sub-structure can be seen inside the breed, some animals had a more distinct cluster and other animals, found in Midwest and Southwest, demonstrated a high degree of admixture, consistent with the previous hypothesis.

Santa Inês animals are an important source of diversity of Brazilian hair sheep with the lowest inbreeding coefficient. Therefore, animal breeding programs focused on different breeding goals are likely to succeed within this breed. The exploitation of the different ecotypes for each environment is a great opportunity for Brazilian sheep industry.

## Conclusions

European and African ancestry was identified for Brazilian hair breeds. Brazilian Somali breed represents an East African group within the Brazilian hair breeds. Brazilian Blackbelly had a clear relationship with Barbados BlackBelly. Brazilian Fat-tail is close to Morada Nova and shows contributions from Brazilian Somali and Afrikaner breeds. Morada Nova has a unique genetic structure. Santa Inês animals came from an admixed population and recently received introgression from Suffolk animals. This breed has a great diversity awaiting for genetic endeavors. The genetic conservation efforts for these breeds is essential due the specific traits and genes that these animals may carry, which would be useful to face further environmental constraints to commercial production. The Brazilian Somali and Blackbelly should receive special attention for conservation efforts.

## Acknowledgments

To CAPES for providing the scholarship for the first author. To CNPq for providing financial support. To Instituto Federal de Educação, Ciência e Tecnologia Goiano for supporting the first author.

## Supporting information

**S1 Table. Description of the 94 breeds used with number of animals (n), Country and region of origin, classification of body cover, tail and purpose.** The breeds used in each principal components analyses are shown with an asterisk (*).

**S2 Table. Results of threepop (f_3_) test performed with the 29 breeds used in Treemix analysis.** The spreadsheet named 100 represent the estimates realized with 100 SNPs per block and the other named 500 represent the estimates with 500 SNPs per block. f_3_: f_3_ statistics; SE: standard error; Significance: means z-score lower than −2 according to Reich et al. (2009); Relationship: indicates the meaning of the analysis using A, B and C as identification of the breeds.

**S3 Table. Results of fourpop (f_4_) test performed with the 29 breeds used in Treemix analysis.** The spreadsheet named 100 represent the estimates realized with 100 SNPs per block and the other named 500 represent the estimates with 500 SNPs per block. f_4_: f_4_ statistics; SE: standard error; Significance: means z-score lower than −2 or higher than 2 according to Reich et al. (2009); Relationship: indicates the meaning of the analysis using A, B, C and D as identification of the breeds, <->: means the most probable relationship in the specific comparison.

**S4 Table. Results of threepop (f_3_) test performed with the 8 breeds used in the second Treemix analysis to refine the genetic background of Brazilian breeds.** The spreadsheet named 100 represent the estimates realized with 100 SNPs per block and the other named 500 represent the estimates with 500 SNPs per block. f_3_: f_3_ statistics; SE: standard error; Significance: means z-score lower than −2 according to Reich et al. (2009); Relationship: indicates the meaning of the analysis using A, B and C as identification of the breeds.

**S5 Table. Results of fourpop (f_4_) test performed with the 8 breeds used in the second Treemix analysis to refine the genetic background of Brazilian breeds.** The spreadsheet named 100 represent the estimates realized with 100 SNPs per block and the other named 500 represent the estimates with 500 SNPs per block. f_4_: f_4_ statistics; SE: standard error; Significance: means z-score lower than −2 or higher than 2 according to Reich et al. (2009); Relationship: indicates the meaning of the analysis using A, B, C and D as identification of the breeds, <->: means the most probable relationship in the specific comparison.

**S1 Fig. Principal component analysis (PCA) using genomic data of 94 breeds from different continents worldwide (according to region in S1 Table).** Blue circle showing the first breeds removed from the analysis. Red circle showing the second group of breeds excluded from posterior analysis. The number between parentheses in each axes means the amount of the variance explained by each eigenvector (EV).

**S2 Fig. Plot of the variance explained by ten eigenvalues in each of the three PCA analyses conducted.** First PCA used data of 94 breeds from different continents worldwide. Second PCA used data of 50 breeds from different continents worldwide and Third PCA used data of 28 breeds genetically close to Brazilian hair sheep breeds.

**S3 Fig. Principal component analysis (PCA) using genomic data of 50 breeds from different continents worldwide (according to region in S1 Table)**. Red circle showing the group of breeds excluded for posterior analysis. The number between parentheses in each axes means the amount of the variance explained by each eigenvector (EV).

**S4 Fig. Plot of the first three eigenvalues of the Principal Component analysis (PCA) using genomic data of 28 breeds genetically close to Brazilian hair sheep breeds**. DOR: African Dorper; AWD: African White Dorper; BBB: Barbados BlackBelly; BAR: BarbarineAlg; BBN: Brazilian Blackbelly; BER: BerberAlg; BBA: Bergamasca; CHI: Chios; COM: Comisana; DMA: DmenAlg; EMZ: Ethiopian Menz; HAM: HamraAlg; MCM: Macarthur Merino; BMN: Morada Nova; NA: Namaqua Afrikaner; ODA: O.DjellalAlg; BPT: Pantaneira; BRL: Brazilian Fat-tail; RM: Red Maasai; REM: RembiAlg; RA: Ronderib Afrikaner; SAK: Sakiz; BSI: Santa Inês; SID: SidaounAlg; BSO: Brazilian Somali; STE: St Elizabeth; SUF: Suffolk; TAZ: TazgezawthAlg.

**S5 Fig. Plot of prediction error of each 10 repetitions for each K value estimated in ADMIXTURE software for 28 sheep breeds data set.** The lowest mean of cross-validation error was obtained for K = 18.

**S6 Fig. Clustering performed with ADMIXTURE software on genotypic data from 28 sheep breeds (K, number of clusters, from 17 to 30).**

**S7 Fig. Plot of prediction error of each 10 repetitions for each K value estimated in ADMIXTURE software for 7 sheep breeds genetically close to Brazilian breeds in the data set.** The best prediction model was considered as K = 4.

**S8 Fig. Inferred trees from 29 sheep breeds using Treemix software simulating 5 and 10 migration events (m = 5 and m = 10, respectively) and using Soay breed as root.** Brazilian breeds highlighted in green.

**S9 Fig. Results of ln(likelihood) for tree simulated with different migration (M) events (0, 1, 2, 3, 4, 5, 10, 20, 30, 40 and 50).**

**S10 Fig. Plot of residual values for each pair of population in the inferred trees with 29 breeds using Treemix software simulating 4, 5, 10 and 20 migration (m) events.** Extreme high residual value means that the representation of that pair of populations in the respective tree underestimates the observed covariance between that populations and extreme negative values means that the representation of that pair of population is overestimating the covariance.

**S11 Fig. Results of ln(likelihood) for tree simulated using 8 breeds (Soay as root) with different migration (M) events (0 to 10) and different size of SNP blocks (1, 100 and 500).**

**S12 Fig. Plot of residual values for each pair of population in the inferred trees with 8 breeds using Treemix software simulating 3 migration events.** Extreme high residual value means that the representation of that pair of populations in the respective tree underestimates the observed covariance between that populations and extreme negative values means that the representation of that pair of population is overestimating the covariance.

